# Using social media to quantify spatial and temporal dynamics of nature-based recreational activities

**DOI:** 10.1101/093112

**Authors:** Francesca Mancini, George M. Coghill, David Lusseau

## Abstract

Big data offer a great opportunity for nature-based recreation (NbR) mapping and evaluation. However, it is important to determine when and how it is appropriate to use this resource. We used Scotland as a case study to validate the use of data from Flickr as an indicator of NbR on a national scale and at several regional spatial and temporal resolutions. We compared Flickr photographs to visitor statistics in the Cairngorms National Park (CNP) and determined whether temporal variability in photo counts could be explained by known annual estimates of CNP visitor numbers. We then used a unique recent national survey of nature recreation in Scotland to determine whether the spatial distribution of Flickr photos could be explained by known spatial variability in nature use. Following this validation work, we used Flickr data to identify hotspots of wildlife watching in Scotland and investigated how they changed between 2005 and 2015.We found that spatial and temporal patterns in Flickr count are explained by measures of visitation obtained through surveys and that this relationship is reliable down to a 10 Km scale resolution. Our findings have implications for planning and management of NbR as they suggest that photographs uploaded on Flickr reflect patterns of NbR at spatial and temporal scales that are relevant for ecosystem management.

## Introduction

Nature-based recreation (NbR) is a key cultural ecosystem service provided by nature and it represents a big component of global recreation [1]. NbR is an important issue because of its economic contribution to conservation [2,3], the health benefits it brings to humans [4] and its role in alleviating poverty [5]. Wildlife watching is a very popular NbR activity that was initially welcomed by conservation and environmental organisations as an eco-friendly alternative to consumptive activities, such as hunting and fishing [6]. However, there is growing evidence that wildlife watching, if not managed properly, can have negative effects on the environment [7–9]. Quantifying temporal and spatial patterns of wildlife watching can help management by identifying areas that are under high pressure from these activities and areas that could be sustainably developed to redistribute this pressure, for example through de-marketing and environmental engineering [10,11].

The widespread use of the Internet and the popularity of smartphones and social media websites offer the opportunity to use the data generated by their billions of users [12,13]. Geotagged information from mobile phones and social media have been used, for example, to estimate the size of crowds [14], a country’s economic activity [15], telecommunication flows [16] and human mobility patterns [17] including international travel flows [18]. A few studies have compared data from social media to visitor statistics obtained through more traditional methods (e.g. surveys or censuses) [19–23]. All of these studies have demonstrated that there is a correlation between these two types of data. Wood et al. [19] and Levin et al. [20] tested the use of social media to quantify NbR in protected areas and selected tourist sites across the globe; Levin et al. [23] and Sessions et al. [21] tested the same on a national scale, respectively in Australia and the United States; while Hausmann et al. [22] validated the use of the content of photographs posted online to infer tourists’ preferences inside the Kruger National Park. Some studies looked at the temporal component of social media data, either explicitly testing whether temporal trends of photographs taken inside national parks and uploaded on Flickr had a robust statistical relationship with visitor statistics [21], or showing that photographs from Flickr were able to idenitfy peaks in visitation to national parks or visitor attractions during popular events [19]. These studies have focused only on visits to protected areas and selected visitor attractions, which are mainly a measure of tourism (domestic and international), while other recreational activities that are carried out in the wider countryside or in urban green networks have not been considered. Moreover, none of them addressed the question of whether the spatial patterns of geotagged photographs from social media correspond to those found in data collected using conventional methods. Therefore, we still do not know if this data can quantify cultural ecosystem services outside protected areas, at which spatial resolution it is accurate and whether we can use it to infer short- and long-term temporal patterns of NbR.

However, following promising results from the first study [19], the use of data from social media to quantify recreational ecosystem services and tourists’ preferences has become increasingly popular [24–32]. Here we make use of two unique independently obtained datasets, resulting from classical survey methodologies, to determine whether temporal and spatial variability in social media photographs are associated with actual spatial and temporal variability in nature recreation. First, we assessed whether the temporal patterns of photographs posted on social media were associated to temporal patterns of visitation to an NbR site and whether changes in the popularity of the social media affected the reliability of its data. We then determined whether areas that are intensely photographed by social media users correspond to areas that are highly visited according to visitor surveys and identified the spatial resolution at which social media photo density is no longer explained by known visitation patterns. Following on this, we created visualisations of photographs of wildlife taken in Scotland and uploaded on Flickr to look for spatio-temporal patterns in wildlife watching activities in Scotland.

## Materials and methods

This paper presents a rare case study in which the patterns observed in social media data were validated against survey data over time and at a relatively fine spatial scale. Wildlife watching contributes roughly £127 million per year to the Scottish economy [33], generating 2763 full-time equivalent jobs [34]. The main groups of charismatic wildlife that attract tourists to Scotland are birds, seals, whales and dolphins [35]. We tested whether temporal patterns of photographs taken at an NbR site (the Cairngorms National Park - CNP) and posted on Flickr corresponded to patterns found in time series of visitor numbers obtained from the CNP authority. Since changes in the popularity of Flickr could affect the reliability of Flickr photographs as a measure of temporal patterns of visitation, we also tested whether the number of active users on Flickr over time had an effect on the time series of Flickr photographs taken in the CNP. We then tested the correspondence between the spatial distribution of geotagged photographs of wildlife posted on Flickr and spatial patterns of wildlife watching activities in Scotland using a dataset collected through a Scotland-wide recreation and tourism survey [36]. Finally, we conducted a "real-world" test of the usefulness of Flickr data as an indicator for wildlife watching activities in Scotland. We looked at the spatiotemporal patterns that emerged from the Flickr photographs and qualitatively determined whether they were reproducing realistic trends in wildlife watching in Scotland.

All the data used in this study are publicly available and were collected following terms and conditions of the data providers. The data are anonymised and we did not collect any personal information. All the R code necessary to reproduce this study is available on GitHub. See Data Accessibility statement for the links to the repositories.

## Data collection

We downloaded data from the Flickr Application Programming Interface (API) to create two datasets. The first dataset was a monthly time series of Flickr photographs taken in the CNP between 2009 and 2014 and it was compared to time series of visitor numbers provided by the CNP authority and to a time series of popularity of Flickr. The second dataset contained the number of Flickr photographs depicting wildlife that were taken on the coast of Scotland between October 2014 and October 2015 and it was compared to a spatial dataset of wildlife watching from the Scottish marine recreation and tourism survey [36]. In the next two sections we describe these datasets before providing details on the analysis.

### Datasets for temporal comparison in nature recreation between social media and survey data

<u>CNP STEAM visitor days</u> - The CNP visitation dataset contained the number of visitor days to the park (a person spending at least a portion of a day at the CNP) per month from 2009 to 2014. These visitor days were estimated by the STEAM model [37], which uses data on accommodation occupancy rates, visitor surveys, number of visitors to paid attractions and other data supplied by the CNP authority. STEAM is not designed to provide a precise and accurate measurement of tourism in a local area, but to provide an indicative base for monitoring trends, and, although it is not a statistically robust measure of tourism, it is one of the most used commercial models in the tourism sector in the UK [37].

<u>CNP Flickr visitor days</u> - Data from Flickr were collected through the Flickr API [38] and R [39], using the packages RCurl [40] version 1.95.4.7, XML [41] version 3.98.1.3 and httr [42] version 1.1.0 to communicate with the API, request and download the data. Dates and geographic coordinates associated with the photographs were used to select only those taken in the CNP between 2009 and 2014. A bounding box was used to query the Flickr API and then a polygon shapefile of the CNP (available at https://data.gov.uk) was used to select only the photographs taken inside the boundaries of the park. We downloaded the following metadata associated with the photographs: photograph and user ID, the date when the photograph was taken and the geographic coordinates of where it was taken. To avoid bias coming from having a small number of very active users, we used the combination of user ID and date to delete multiple photographs from the same user on the same day, thus retaining only the first photograph taken every day by each user. By counting the number of photographs retained in each month we then obtained the monthly number of Flickr visitor days in the CNP (a person taking at least one photograph a day in the CNP).

<u>Flickr active users</u> - To quantify changes in the popularity of Flickr over the years, we used the number of active users (i.e. users posting content on Flickr). We downloaded a publicly available global dataset of 100 million media objects uploaded on Flickr from 2004 to April 2014 [43] and calculated Flickr active users as the number of users who posted at least one media object on Flickr in each month from January 2009 to April 2014 (Figure A in S1 Appendix).

### Datasets for spatial comparison in wildlife watching activity between social media and survey data

<u>Coastal survey visitor days</u> - The dataset from the Scottish marine recreation and tourism survey [36] contained spatial information on trips to coastal areas in Scotland made by the survey respondents between October 2014 and October 2015 to conduct wildlife watching activities. The online survey asked respondents to draw polygons on a map of Scotland, corresponding to the areas they had used for wildlife watching activities in that period. All the polygons were then stacked on top of each other and counted, resulting in a raster file with a resolution of 1 km.

This dataset, as most survey data, has some limitations and weaknesses. The survey was sent to a number of relevant organisations (such as clubs, tour operators, public agencies etc.), advertised through press releases to key media organisations, user groups, industry organisations and on social media (Facebook and Twitter). As a result of this dissemination strategy, the survey was not based on a random sample of the Scottish population, therefore it is likely to have a bias towards a proportion of the population that is highly active in outdoor coastal activities. A total of 933 people provided spatial information on the places they visited to take part in wildlife watching activities, however, the data is not likely to be fully comprehensive. There was a small number of responses from remote parts of Scotland, which means that the spatial information for areas such as the Western Isles and Shetland is likely to be partial.

<u>Coastal Flickr visitor days</u> - We used keywords (bird, seal, dolphin and whale) to query the Flickr API and select all the geotagged photographs of the main groups of wildlife sought by wildlife watchers in Scotland and taken between October 2014 and October 2015. We spatially restricted the query to the Flickr API by providing Scotland’s WOEID (Where On Earth IDentifier), which is a unique 32-bit reference identifier assigned by Yahoo! that identifies any feature on Earth. We downloaded the same metadata as for the CNP Flickr visitor days, but, in addition, we downloaded the user tags, which we used to eliminate photographs that were not relevant (such as photographs of statues or paintings and photographs taken in zoos). The tags were examined and a list of keywords for nonrelevant photographs was compiled, then following a method similar to that used in [32], we used that list of keywords to filter out irrelevant photographs. We recognise that this method is not perfect and it might have retained some non-relevant photographs, but we suggest that, considering the high number of photographs we processed, this was the best way to ensure that the mistakes introduced by the filtering procedures were not biasing the results. A buffer of 2 km inside the coastline was created in ArcGIS [44] to select only the photographs taken in a coastal environment. For each of the different wildlife groups, we again deleted multiple photographs from the same user on the same day. The exact location of each photograph is important in this part of the analysis and it is possible that by only keeping the first photograph by each user every day we have removed some photographs that were taken in a different location from the first one. Therefore, we calculated the distance between multiple photographs taken by the same user on the same day. Since the median distance is 0 m and the mean is 3 Km, we believe that this filtering procedure did not significantly affect the results (Figure B in S1 Appendix). Finally, we created 3 grids (5 km, 10 km and 20 km) and counted the number of Coastal Flickr visitor days and of Coastal survey visitor days in each cell.

## Analysis

### Temporal comparison in nature recreation between social media and survey data

In order to test whether the temporal patterns shown by the CNP Flickr visitor days and the CNP STEAM visitor days time series were similar, we used Wavelets Analysis (WA) [45]. WA decomposes the variance of the time series in its oscillating components, thus detecting significant periodicities. The advantage of this method compared to other spectral decompositions is that WA does not assume stationarity of the time series, but it allows the main frequency component to change through time by estimating the signal’s spectral characteristics as a function of time (the wavelet power spectra). We used the Morlet mother wavelet to perform this decomposition [46]. This continuous and complex function allows the extraction of both time-dependent amplitude and phase of the time series. WA also allows the analysis of patterns of covariation between two time series [47–50]. We compared the time series of CNP STEAM visitor days and CNP Flickr visitor days using the wavelet coherence, which identifies the linear correlation between two signals. In order to assess statistical significance of the association between the two time series we used a random noise resampling scheme, where the null hypothesis tested is that the association between the two signals is not different from that expected by chance alone [47]. We also computed the phase difference to test whether the two time series were synchronised or out of phase. This analysis was conducted in R using the package WaveletComp version 1.0 [51].

To exclude the possibility that the time series of CNP Flickr visitor days was influenced by the changes in the popularity of Flickr, we tested whether the number of Flickr active users had an effect on CNP Flickr visitor days. We fitted three Generalised Estimating Equations (GEE) with autoregressive correlation structures and the following formulas:

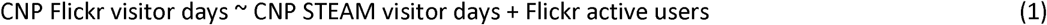

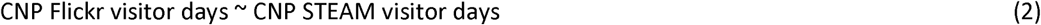

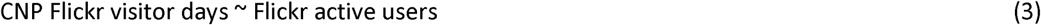

Due to the strong seasonal trend in both the CNP STEAM visitor days and the Flickr active users, we checked for collinearity between these two variables. We used the function vifcor in the R package usdm [52]. Given a variance inflation factor (VIF) < 3 we determined that we could use both variables in the same model [53]. We fitted the GEE models in R [39] using the package geepack [54–56]. We then compared these three models using quasi-likelihood under the independence model criterion (QIC).

### Spatial comparison in wildlife watching activity between social media and survey data

We compared the spatial distribution of wildlife watchers obtained from Flickr (Coastal Flickr visitor days) and the one obtained from the survey (Coastal survey visitor days) at three different spatial scales: 5 km, 10 km and 20 km. We fitted Generalised Linear Models (GLM) to the three datasets using the number of Coastal Flickr visitor days in each cell as the response variable and the number of Coastal survey visitor days in each cell as explanatory variable. The data contained a high number of zeros, so we fitted a GLM with a binomial error distribution to the presence/absence of Coastal Flickr visitor days in a cell and a GLM with a negative binomial error distribution to the number of Coastal Flickr visitor days present. This allowed us to test two hypotheses: 1) the probability of finding at least one photograph by one user on Flickr is higher for areas with higher Coastal survey visitor days 2) the number of users posting photographs on Flickr is higher for areas with higher Coastal survey visitor days. Since densely populated areas tend to have a higher average number of Flickr users (Figure C in S1 Appendix), we used population abundance for each grid cell as model weights. The data for population size was a 1 km resolution raster of estimates of population count available at [57]. The residuals of the GLMs were spatially correlated and directional variograms, estimated using R package gstat version 1.1.0 [58], showed that the spatial autocorrelation was anisotropic (Figure D in S1 Appendix). We therefore used spatial eigenvector mapping (SEVM) to derive explanatory variables for the GLMs [59]. This method decomposes a matrix of relationships between data points into eigenvectors that capture spatial effects. The eigenvectors can then be included as explanatory variables in the GLM to remove the effect of spatial autocorrelation on the analysis. First we used a Delaunay triangulation of the grid cells centres to define neighbours, to which we assigned row-standardised spatial weights. A set of Moran eigenvectors (ME) were then calculated from these weights and those that best reduced the spatial autocorrelation of residuals were selected and included as spatial covariates in the GLMs. We used AIC to select only the ME that improved the explanatory power of the model. This analysis was conducted in R using the package spdep version 0.5.92 [60,61].

## Results

In total, we downloaded metadata on 29,336 photographs (4699 unique CNP Flickr visitor days) taken in the CNP between 2009 and 2014 and uploaded on Flickr.

The query to Flickr API returned 92,229 results (36,998 Flickr visitor days and 2,658 Coastal Flickr visitor days) for photographs with the word “bird” in tags, title or description taken in Scotland between 2005 and 2015. From the search with the word “seal” we obtained 7212 photographs (242 Coastal Flickr visitor days) and from the search with the word “dolphin” and the word “whale” we obtained 5994 photographs (94 Coastal Flickr visitor days).

### Temporal comparison in nature recreation between social media and survey data

The power spectra of the Flickr and CNP STEAM time series were very similar, with significant 12- month cycles throughout the 5-year period (Figure E in S1 Appendix), showing that visitation has a strong seasonal component in both measures. This similarity was supported by the strong coherence between the Flickr and the empirical time series around the same 12-month oscillations, which was also constant through time (Fig 1a). The phase difference for this 12-month cycle was constant around 0 (Fig 1b), indicating that the two time series were synchronised. Cross-correlation was also significant at a period of 6 and 3 months, but this was not consistent throughout the time period (Fig 1a).

**Fig 1.**
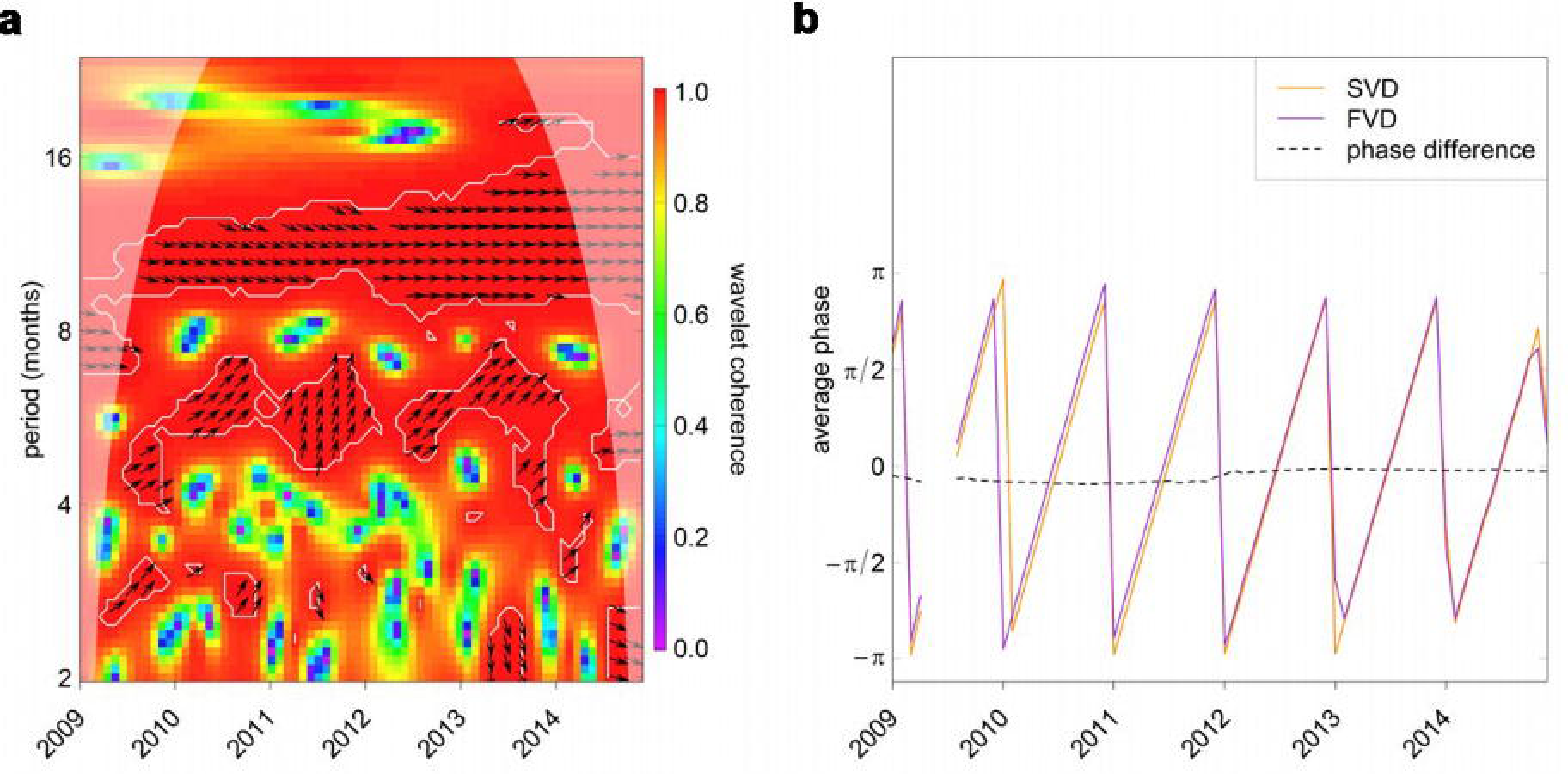
Results of wavelet analysis. a) Wavelet coherence between the two time series. Colour code from purple (low values) to red (high values). The arrows indicate synchrony of the two time series: arrows pointing to the right means the oscillations are synchronised. Arrows are only plotted within white contour lines that indicate significance. The shaded area near the edges in the graphs is the cone of influence, and indicates the range of the graph where the results are not reliable because of edge effects, b) Phases of the oscillations of the two time series (SVD - STEAM visitor days - in orange and FVD - Flickr visitor days - in purple) computed in the 8-16 periodic band where there is significant correlation. The dotted line is the phase difference.

The best model according to QIC was the first model including both CNP STEAM visitor days and Flickr active users (Table A in S1 Appendix). Excluding the variable CNP STEAM visitor days resulted in a jump of 334.75 in QIC (Table A in S1 Appendix), indicating that this variable was explaining most of the variability in CNP Flickr visitor days. We could not detect a significant effect of the number of Flickr active users on the time series of CNP Flickr visitor days (Wald = 0.043; SE = 7.599e-06; p > 0.05), while CNP STEAM visitor days explained a significant proportion of the variability in the response variable (Wald = 32.129; SE = 3.823e-07; p < 0.001).

### Spatial comparison in wildlife watching activity between social media and survey data

The Coastal survey visitor days dataset indicated that the areas more intensely used for wildlife watching are around the west coast of Scotland, the Moray Firth, the Firth of Forth and the Tay estuary (Figure F in S1 Appendix). Spatial distribution of the photographs from Flickr also identified the last three as areas of high visitation (Figure C in S1 Appendix). The Coastal Flickr visitor days dataset also contained photographs taken on the west coast of Scotland, but not in the same density as shown by the survey dataset. This area of the country is not highly populated, so there might be a certain bias in the number of Flickr users uploading photographs. When the Flickr data was normalised by population size, the west coast appeared as an area of high visitation (Figure G in S1 Appendix).

The probability of finding a Flickr photograph taken in a certain cell was higher in cells with higher Coastal survey visitor days (Fig 2). This was true for each of the spatial scales tested (20 km: estimate = 0.37, *SE* = 0.04, *Z* = 9.5, *p*-value < 0.001; 10 km: estimate = 0.36, SE = 0.02, Z = 14.7, *p-value* < 0.001; 5 km: estimate = 0.28, *SE* = 0.001, *Z* = 21.1, *p-value* < 0.001). The higher the number of visitors captured by the survey, the higher the probability of finding photographs on Flickr taken in that area.

**Fig 2.**
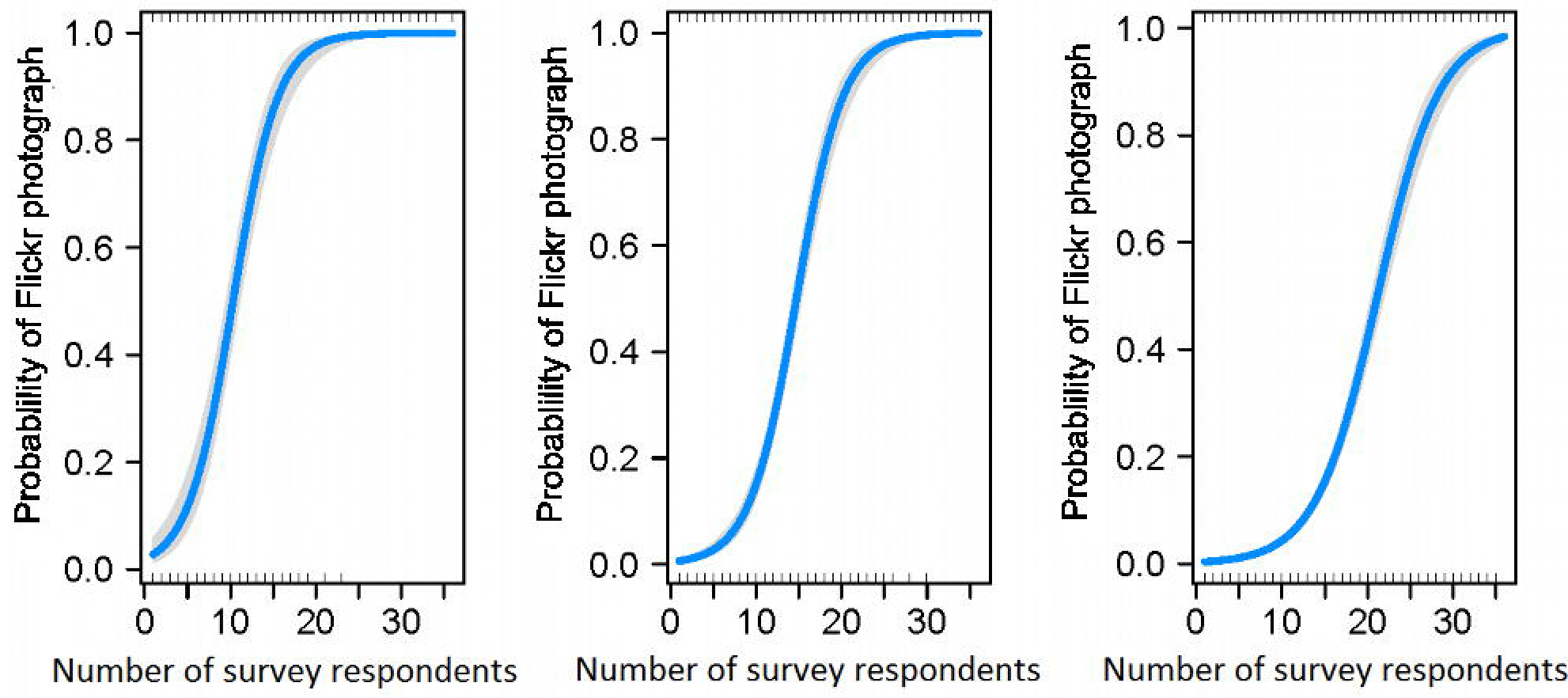
Results of binomial GLMs. Left: results at the 20 km resolution; centre: results at the 10 km resolution; right: results at the 5 km resolution. Predictions from the models (blue line) are plotted on the response scale with confidence intervals (shaded areas around the prediction curve).

We found a significant correspondence between the number of Coastal Flickr visitor days and the number of Coastal survey visitor days at 10 km and 20 km resolution but not at 5 km (Fig 3; 20 km: estimate = 0.1, *SE* = 0.01, *Z* = 8.1, *p-value* < 0.001; 10 km: estimate = 0.02, *SE* = 0.001, *Z* = 3.4, *p-value* < 0.001; 5 km: estimate = 0.01, *SE* = 0.006, *Z* = 1.7, *p-value* > 0.05). The number of Flickr users taking a photograph in a cell was significantly higher in cells with higher number of visitors captured by the survey, but only at a 10 and 20 km resolution.

**Fig 3.**
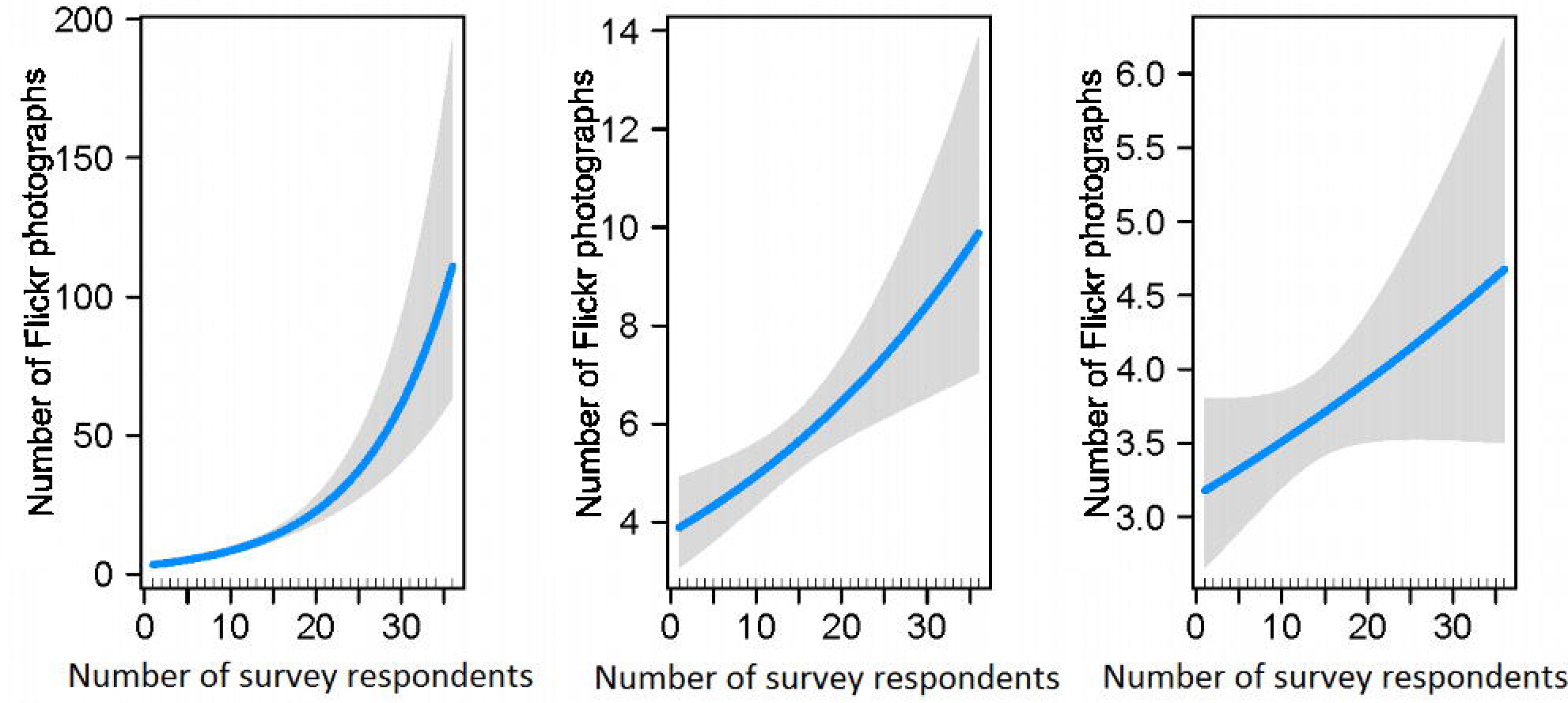
Results of negative binomial GLMs. Left: results at the 20 km resolution; centre: results at the 10 km resolution; right: results at the 5 km resolution. Predictions from the models (blue line) are plotted on the response scale with confidence intervals (shaded areas around the prediction curve).

### Visualising spatio-temporal patterns of wildlife watching activities in Scotland

We collected a third dataset containing metadata of photographs of birds, seals, whales and dolphins taken in Scotland from 2005 to 2015 and uploaded on Flickr, using the same methodology described in the Methods section. We visualised spatio-temporal patterns in wildlife watching hotspots by producing density maps of the geotagged photographs. Our aim was to show what Flickr photographs can tell us about wildlife watching in Scotland and determine whether the trends emerging from the data corresponded to known patterns of wildlife watching in Scotland. We used a two-dimensional kernel density estimator from the R package ggplot2 [62], where the bandwidth is calculated using Scott’s rule of thumb [63].

The density maps revealed spatio-temporal patterns of wildlife watching hotspots in Scotland. Bird watching (Fig 4) was concentrated around Edinburgh and Glasgow. However, season played an important role in determining the distribution of bird watchers, in fact the activity was most concentrated in the urban areas in winter and spread out towards the West coast and the islands during spring and summer (Fig 4).

**Fig 4.**
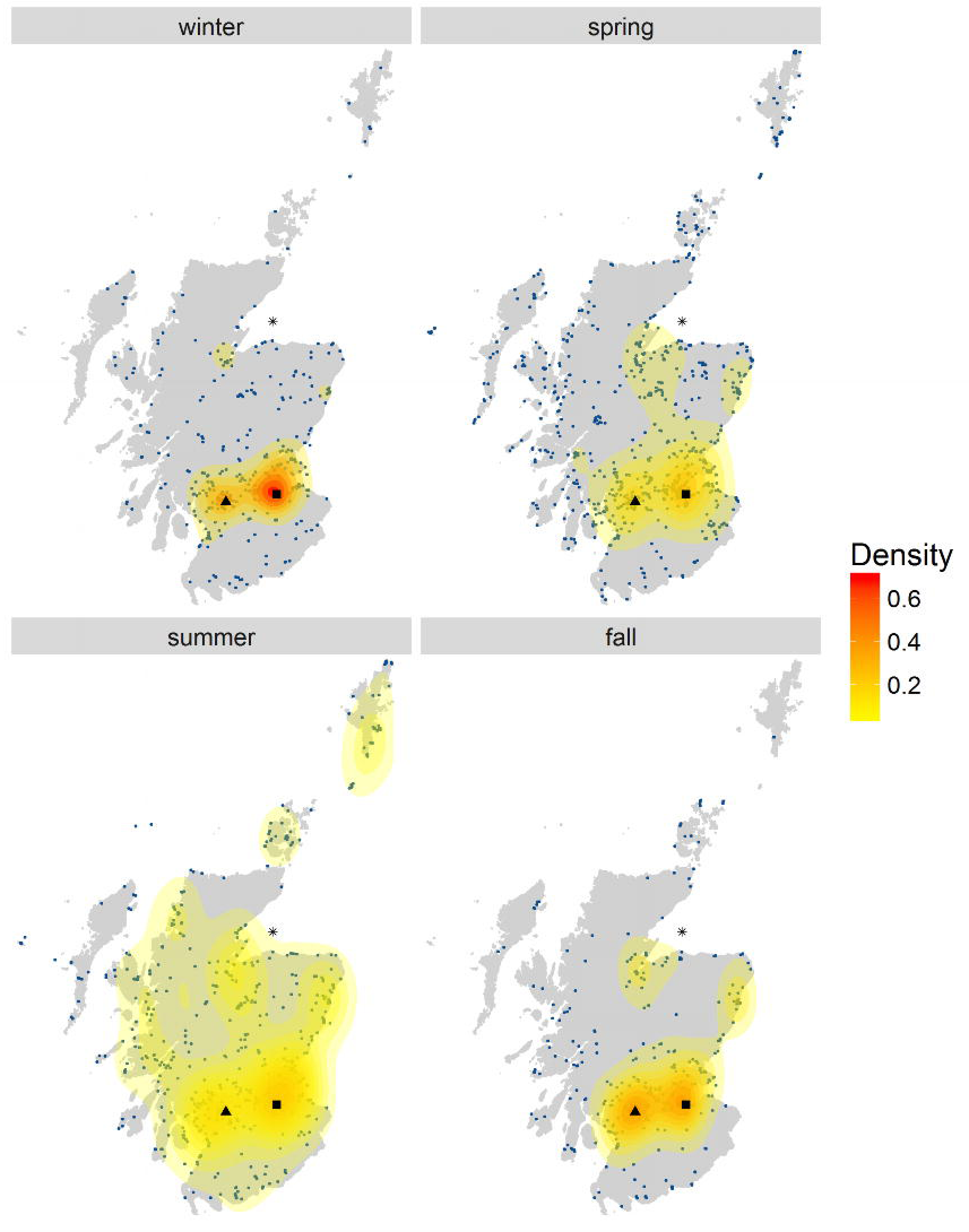
Birdwatching density map for 2009 by season. Each panel represents the density of Flickr visitor days in a different season. The blue dots on the maps are the data. Different colours represent different density levels, from low (yellow) to high (red). * Moray Firth, ͳ Edinburgh; ▲ Glasgow.

Seal watching was concentrated initially around the west coast, the Firth of Forth and Shetland (Fig 5). It is worth noticing the appearance of another hotspot from 2008 corresponding to Newburgh in Aberdeenshire, becoming very popular after 2011.

**Fig 5.**
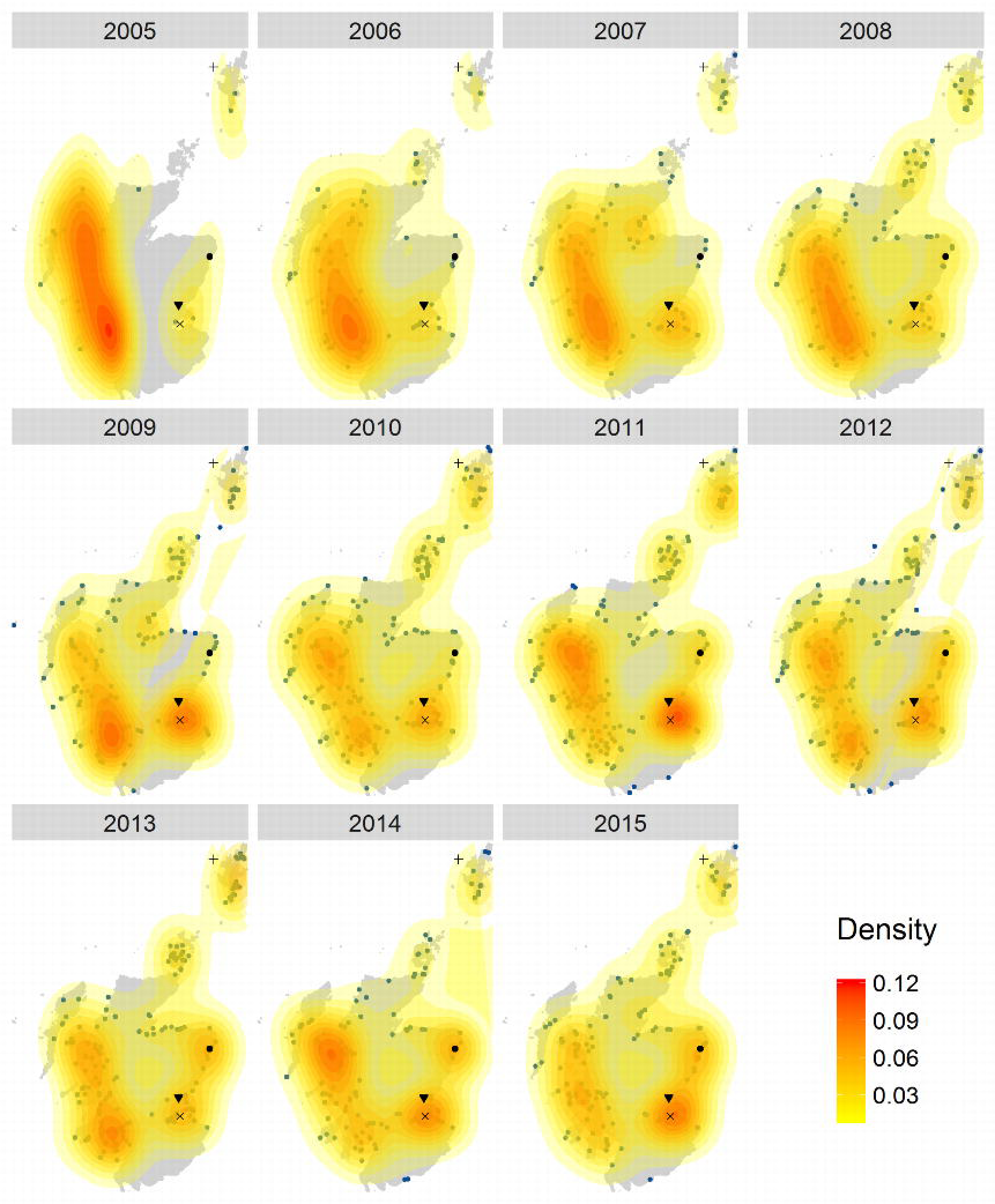
Seal watching density maps. Each panel represents the density of Flickr visitor days in a different year, from 2005 to 2015. The blue dots on the maps are the data. Different colours represent different density levels, from low (yellow) to high (red). + Shetland; ● Newburgh; ▼ Tay estuary; x Firth of Forth.

Dolphin and whale watching maps showed a consistent hotspot at Chanonry Point in the Moray Firth (Fig 6), with the appearance of a second hotspot from 2013 in Aberdeen.

**Fig 6.**
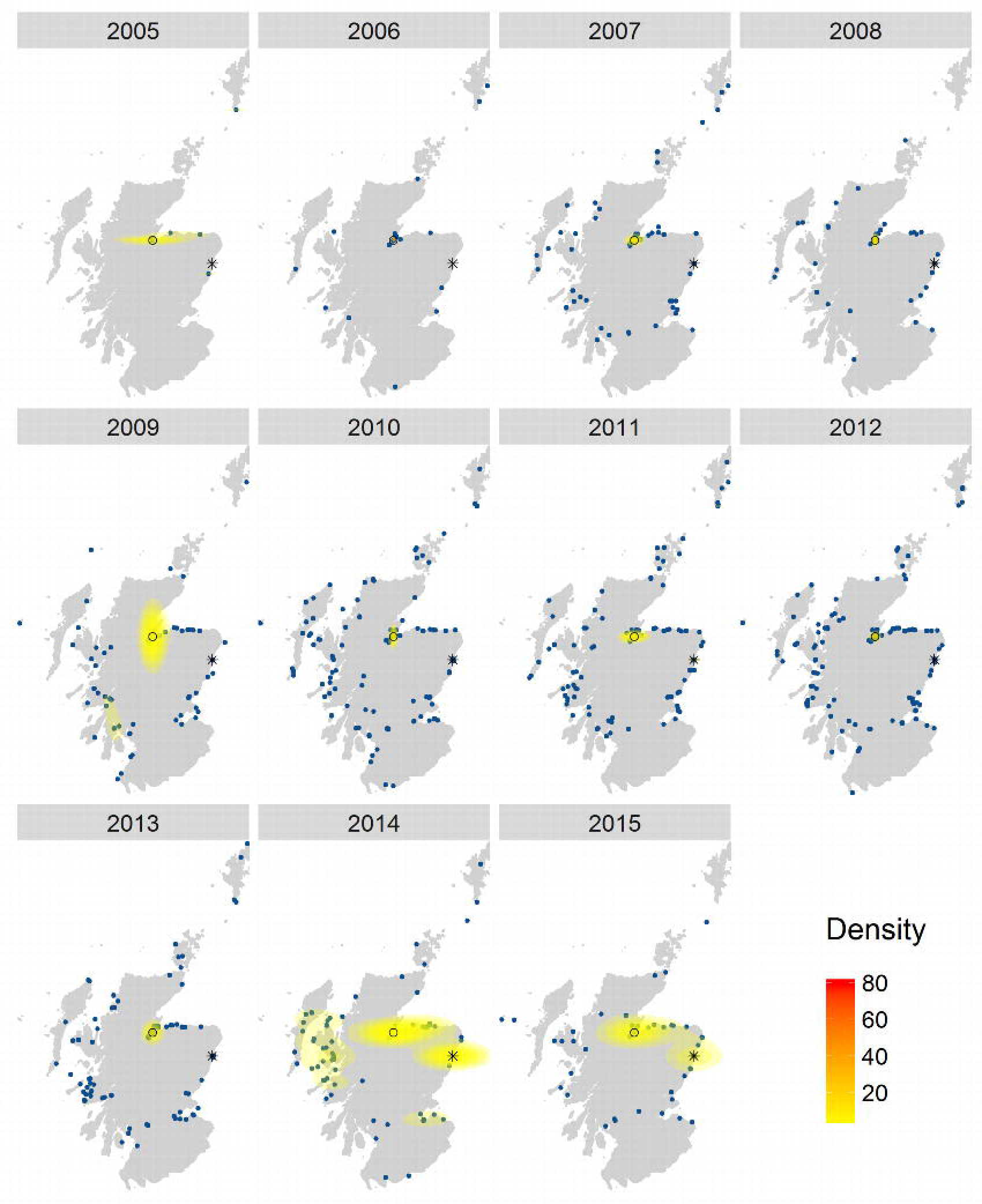
Dolphin and whale watching density maps. Each panel represents the density of Flickr visitor days in a different year, from 2005 to 2015. The blue dots on the maps are the data. Different colours represent different density levels, from low (yellow) to high (red). ○ Chanonry Point; * Aberdeen.

## Discussion

In this study, we were able to compare spatial and temporal patterns in photographs from Flickr to those obtained through traditional methods. The results are in line with previous work [19–23], suggesting that geotagged photographs uploaded on Flickr reflect patterns of NbR activities. Novel results from our study reveal that there is a reliable statistical relationship between spatial patterns of Flickr photographs and of survey data on a national scale at a resolution as fine as 10 Km (Figs 2 and 3) and that changes in the popularity of Flickr do not affect this relationship in the study period.

### Temporal comparison in wildlife-watching activity between social media and survey data

Photographs uploaded on Flickr show the same seasonal oscillations found in visitor statistics for an NbR destination. Despite the strong correlation between the seasonal oscillations of the two time series, annual trends showed some differences (Fig 7). These changes could not be explained by changes in the popularity of Flickr as we found no significant effect of the number of Flickr active users on the time series of CNP photographs.

**Fig 7.**
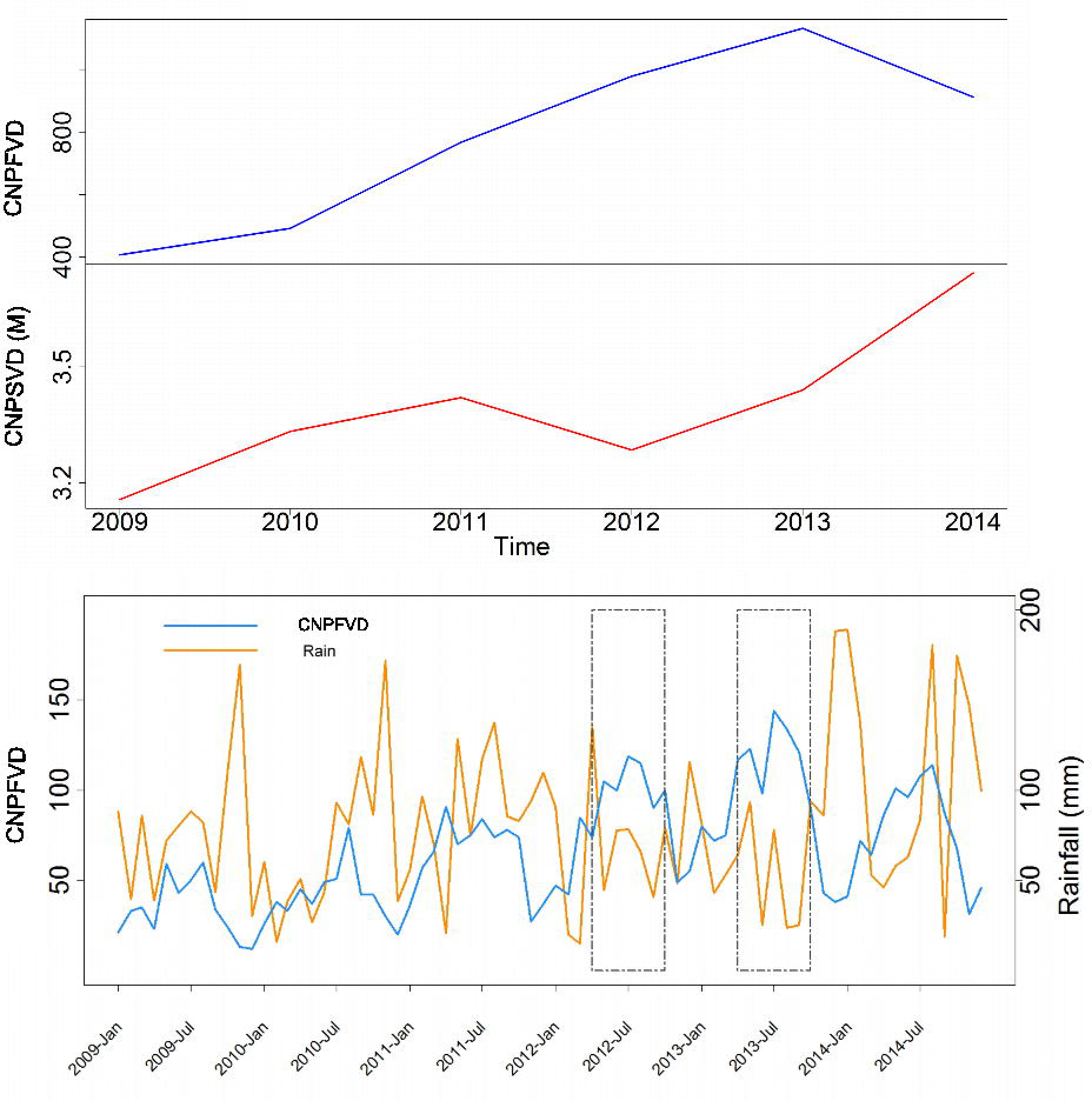
Time series of visitation to the CNP. Top panel: time series of annual visitation to the Cairngorms National Park from Flickr (CNPFVD - CNP Flickr visitor days) and from the CNP authority data (CNPSVD - CNP STEAM visitor days in millions). Bottom panel: time series of monthly Flickr visitor days and rainfall in the Cairngorms National Park. The rectangles indicate summer 2012 and 2013 when rainfall is low and visitation is high. Rainfall data available from Met Office UK at: https://www.metoffice.gov.uk/pub/data/weather/uk/climate/stationdata/braemardata.txt

A possible explanation for the discrepancy in the annual trends is that there is a bias in the data modelled by the STEAM model [37]. The STEAM model is not accurate in terms of absolute number of visitors and it relies on accommodation data, number of visitors to paid attractions and data from visitor surveys provided by the CNP authority [64]. The only visitor survey conducted by the CNP authority was in 2009/2010, therefore day trips for years after 2009/2010 were estimated on the basis of this last survey. The biggest discrepancy in annual trends between the two datasets is found between 2012 and 2014 (Fig 7). Weather conditions in these three years could explain these differences. In 2012 and 2013 precipitations in the summer were very low, while 2014 was a wetter year (Fig 7). We know that, differently from long stays, day trips are heavily influenced by weather conditions [65], therefore estimates based on data collected in 2009-2010 might not be representative of actual numbers of day visitors in years with lower precipitations. Further analysis is needed to explain this discrepancy with more certainty.

### Spatial comparison in wildlife-watching activity between social media and survey data

The spatial resolution at which we can use this data source can be as fine as 5 km (Fig 2), but, at a resolution finer than 10 km, the number of Flickr users taking photographs of wildlife may not be a reliable measure for volume of recreation (Fig 3). This could be due to lack of precision in entering geographic coordinates of the location where the photograph was taken. If GPS signal is not available or the photograph is taken with a device that does not record geographic coordinates, then users can add the spatial reference during the uploading of the photograph on the website by indicating on a map the location where the photograph was taken. This process might not be accurate enough to provide reliable information about where people go at a very fine spatial scale. This result calls for caution when using Flickr data to make inferences about spatial patterns of visitation rates on a regional scale and fine resolution. As the use of smartphones and the quality of the GPS on these devices increase, we should see an improvement in the spatial accuracy of the photographs uploaded and, consequently, an increase in the resolution at which the data will be reliable.

### Visualising spatio-temporal patterns of wildlife watching activities in Scotland

Using both spatial and temporal information from geotagged photographs uploaded on Flickr at the same time provided insights into long-term spatio-temporal patterns of wildlife watching activities that further validated the use of this data source and gave information that is relevant for the management of these activities. The majority of photographs of birds were taken around Edinburgh and Glasgow (Figure H in S1 Appendix). However, we also detected a change in the bird watching hotspots with seasons (Fig 4), consistent with a movement from the area around Edinburgh and Glasgow to more remote areas on the west coast, the Moray Firth and the islands. We were therefore able to capture the movement of people towards NbR destinations [34,36].

The seal hotspot maps (Fig 5) revealed high activity in the Firth of Forth, Tay estuary and the West coast where special areas of conservation (SACs) and haul out sites are present for both grey (*Halichoerus grypus*) and harbour seals (*Phoca vitulina*) [66]. This map also showed the appearance of a seal watching point in Newburgh after 2008. This site now holds 26% of grey seals and 1% of harbour seals in the East Coast of Scotland Seal Management Area and has recently been designated as a haul-out site to protect the seals [67,68]. The popularity of this site kept increasing from 2008 through to 2015 and this data could be important to monitor seal watching activities in the future and determine if more protection measures need to be put in place.

The whale and dolphin watching density maps (Fig 6) showed a very consistent hotspot on the East coast, corresponding to Chanonry Point in the Moray Firth, which is a very popular dolphin watching destination [69]. The same maps also revealed the emergence of a new dolphin watching hotspot in Aberdeen. The hotspot only appears in 2013, after the launch by the Royal Society for the Protection of Birds (RSPB) of dolphin watching events from Aberdeen harbour as part of the "Dates with nature" projects [70]. This result indicates that such organised events can attract attention and create a wildlife watching hotspot, offering an opportunity for managers to influence recreational demand. Further analysis could explore whether the birth of a new hotspot creates new demand or whether it could shift wildlife watchers’ attention from overexploited destinations to unused ones.

## Limitations

Data from social media still presents some limitations that need to be acknowledged, some of which also apply to traditional sampling methods [19,71]. There is a bias resulting from densely populated areas having more Flickr users than sparsely populated ones [20] (Fig 4 and Figure A in S1 Appendix). Different species of wildlife may be more or less suited to be photographed, therefore it might not be possible to compare volume of recreational activities dedicated to different species. Furthermore, the perceived value of a trip may influence whether an individual takes or shares photographs, producing a bias against images from visitors who visit areas closer to their home [19]. Half of the respondents to the marine recreation and tourism survey lived within one mile of the coast [36] and they might have reported using an area where they would not normally take photographs because of its proximity to home. This could explain some of the differences between the two datasets.

It is also important to consider shifts in the popularity of Flickr as a possible source of bias. Our analysis indicated that, in our study period (between 2009 and 2014), changes in Flickr popularity did not influence the time series of photographs taken in an NbR destination (Table A in S1 Appendix). However, Flickr’s popularity has been decreasing in the last 3-4 years, while other photo-sharing websites have been growing. This might compromise the reliability of Flickr as an indicator of visitation in the future, while social media such as Instagram will become representative of a wider proportion of the population.

## Opportunities

Despite these limitations, geotagged photographs uploaded on Flickr show patterns that correspond to NbR survey data at spatial and temporal scales that are relevant for ecosystem management. This opens new avenues to study wildlife watchers’ behaviour and decisions. The fact that we can use this data at a scale as fine as 10 km means that we can now make more precise inference on recreationists’ preferences on larger areas. This information has also implications for wildlife recreation management and conservation of targeted species. First, we can now easily and cheaply quantify wildlife watching in areas that are not monitored, allowing us to assess whether some areas are receiving increasing pressure from recreational activities, while other areas are underutilised. Secondly, organised events such as the RSPB "Date with Nature" can attract wildlife watchers and create a recreational hotspot, offering an opportunity for managers to influence demand. NbR is often difficult to incorporate in ecosystem services exploitation management schemes alongside more traditional activities [72] and therefore it is not given priority. We show here that social media is a useful sampling platform to appraise the scope of NbR at relevant spatial and temporal scales. We can use this information to compare the spatial distribution of NbR to that of other ecosystem services to understand where they overlap and what are the synergies and trade-offs. For example it would be possible to use the detailed spatial and temporal information provided by geotagged photographs to test whether NbR is helping achieve national or global biodiversity targets or identify areas where NbR is in conflict with biodiversity conservation. Lastly, data from social media can give us a dynamic view of demand for NbR, which can be used to anticipate changes in visitation in response to changes in ecosystems and socio-economic development. In a fast-changing world, the ability to predict changes is going to be increasingly important to plan for a sustainable use of the natural environment.

## Conclusions

Short-term and seasonal patterns of photographs uploaded on Flickr show a reliable statistical relationship with temporal patterns of visitation to an NbR destination captured by classical data collection methods. In this study we found a strong correspondence between time series of photographs taken in the CNP and time series obtained from surveys. We also couldn’t find a significant effect of changes in the popularity of the social media on the time series of photographs taken in the CNP.

We also found that photographs of wildlife uploaded on Flickr are reliably described by spatial patterns of wildlife recreational activities captured by a survey on a national scale. The spatial resolution at which this statistical relationship exists can be as fine as 5 km (Fig 2), but, at a resolution finer than 10 km, the number of Flickr users taking photographs of wildlife may not be a reliable measure for volume of recreation (Fig 3). Therefore, we should use caution when using this indicator to infer very fine spatial movement of nature-based recreationists.

The spatio-temporal trends from wildlife photographs uploaded on Flickr confirmed the usefulness of this data in real-world settings. The photographs showed realistic patterns of seasonal movement towards touristic areas, some well-known wildlife watching hotspots in Scotland and their change through time due to organised events and marketing.

The use of data from Flickr will provide opportunities for research into visitor’s preferences and behaviour and for monitoring the pressure from recreational activities in areas that are difficult to monitor, thus helping resolve potential conflicts between NbR, biodiversity conservation and other ecosystem services.

## Acknowledgements

We thank the Marine Alliance for Science and Technology for Scotland (MASTS) and Scottish Natural Heritage (SNH) for their support. We also thank B. Leyshon and F. Manson (SNH) for fruitful discussion.

## Supporting information

**SI Appendix. Supplementary maps and figures.**

**Figure A in S1 Appendix. Flickr Active Users.** Time series showing how many users posted at least one picture on Flickr (Flickr active users) in each month from January 2009 to April 2014.

**Figure B in S1 Appendix. Distance between multiple photographs by the same user.** Distribution of the distance (in meters) between multiple photographs taken by each user on the same day.

**Figure C in S1 Appendix. Flickr visitor days.** Spatial distribution of Flickr visitor days (FVD).

**Figure D in S1 Appendix. Variograms.** Directional variograms of residuals of binomial GLM at the 5km resolution. Each panel shows semivariance in residuals in different directions: from North (panel 0) to North West (panel 320).

**Figure E in S1 Appendix. Time series wavelet power spectra.** Wavelet power spectra of the two time series: a) CNP authority and b) Flickr time series. Colour code from dark blue (low values) to red (high values). White contour lines indicate significance. The shaded area near the edges in the graphs is the cone of influence, and indicates the range of the graph where the results are not reliable because of edge effects.

**Figure F in S1 Appendix. Survey visitor days.** Spatial distribution of coastal survey visitor days.

**Figure G in S1 Appendix. Flickr visitor days normalised.** Spatial distribution of coastal Flickr visitor days normalised by population size.

**Figure H in S1 Appendix. Bird watching density maps.** Each panel represents the density of Flickr visitor days in a different year, from 2005 to 2015. The blue dots on the maps are the data. Different colours represent different density levels, from low (yellow) to high (red).

**Table A in S1 Appendix. QIC of the three alternative models.**

